# CODC: A copula based model to identify differential coexpression

**DOI:** 10.1101/725887

**Authors:** Sumanta Ray, Snehalika Lall, Sanghamitra Bandyopadhyay

**Author notes:** these authors contributed equally to this work (Sumanta Ray, Snehalika Lall).

## Abstract

Differential coexpression has recently emerged as a new way to establish a fundamental difference in expression pattern among a group of genes between two populations. Earlier methods used some scoring techniques to detect changes in correlation patterns of a gene pair in two conditions. However, modeling differential coexpression by mean of finding differences in the dependence structure of the gene pair has hitherto not been carried out.

We exploit a copula-based framework to model differential coexpression between gene pair in two different conditions. The Copula is used to model the dependency between expression profiles of a gene pair. For a gene pair, the distance between two joint distributions produced by copula is served as differential coexpression. We used five pan-cancer TCGA RNA-Seq data to evaluate the model which outperforms the existing state-of-the-art. Moreover, the proposed model can detect a mild change in the coexpression pattern across two conditions. For noisy expression data, the proposed method performs well because of the popular scale-invariant property of copula. Additionally, we have identified differentially coexpressed modules by applying hierarchical clustering on the distance matrix. The identified modules are analyzed through Gene Ontology terms and KEGG pathway enrichment analysis.

## Introduction

Microarray based gene coexpression analysis has been demonstrated as an emerging field which offers opportunities to the researcher to discover coregulation pattern among gene expression profiles. Genes with similar transcriptomal expression are more likely to be regulated by the same process. Coexpression analysis seeks to identify genes with similar expression patterns which can be believed to associate with the common biological process^1–3^. Recent approaches are interested to find the differences between coexpression pattern of genes in two different conditions^4, 5^. This is essential to get a more informative picture of the differential regulation pattern of genes under two phenotype conditions. Identifying the difference in coexpression patterns, which is commonly known as differential coexpression is no doubt a challenging task in computational biology. Several computational studies exist for identifying change in gene coexpression patterns across normal and disease states^6, 6–9^. Finding differentially coexpressed (DC) gene pairs, gene clusters, and dysregulated pathways between normal and disease states are most common.^6, 10–13^. Another way for identifying DC gene modules is to find gene cluster in one condition and test whether these clusters show a change in coexpression patterns in another condition significantly.^8, 14^.

For example, CoXpress^10^ utilizes hierarchical clustering to model the relationship between genes. The modules are identified by cutting the dendrogram at some specified level. It used a resampling technique to validate the modules coexpressed in one condition but not in other. Another approach called DiffCoex^11^ utilized a statistical framework to identify DC modules. DiffCoex proposed a score to quantify differential coexpression between gene pairs and transform this into dissimilarity measures to use in clustering. A popularly used tool WGCNA (Weighted Gene Coexpression Network Analysis) is exploited to group genes into DC clusters^15^. Another method called DICER(Differential Correlation in Expression for meta-module Recovery)^16^ also identifies gene sets whose correlation patterns differ between disease and control samples. Dicer not only identifies the differentially coexpressed module but it goes step beyond and identifies meta-modules or class of modules where a significant change in coexpression patterns is observed between modules, while the same patterns exist within each module. In another approach, Ray et al^17^ proposed a multiobjective framework called DiffCoMO to detect differential coexpression between two stages of HIV-1 disease progression. Here, the algorithm operates on two objective functions which simultaneously optimize the distances between two correlation matrices obtained from two microarray data of HIV infected individuals.

Most of the methods proposed some scoring technique to capture the differential coexpression pattern and utilized some searching algorithm to optimize it. Here, we have proposed CODC **Co**pula based model to identify **D**ifferential **C**oexpression of genes under two different conditions. Copula^18, 19^ produces a multivariate probability distribution from multiple uniform marginal distribution. It extensively used in high dimensional data applications. In the proposed method, first, a pairwise dependency between gene expression profile is modeled using an empirical copula. As the marginals are unknown, so we used empirical copula to model the joint distribution between each pair of gene expression profiles. To investigate the difference in coexpression pattern of a gene pair across two conditions, we compute a statistical distance between the joint distributions. We hypothesized that the distance between two joint distributions can model the differential coexpression of a gene pair between two conditions. To investigate this fact we have performed a simulation study that provides the correctness of our method. We have also validated the proposed method by applying it in real life datasets. For this, we have used five pan cancer RNA-Seq data from TCGA: Breast invasive carcinoma [BRCA], Head and Neck squamous carcinoma [HNSC], Liver hepatocellular carcinoma [LIHC], Thyroid carcinoma [THCA] and Lung adenocarcinoma [LUAD] which are publicly available in TCGA data portal (https://tcga-data.nci.nih.gov/docs/publications/tcga/)

## Materials and methods

In this section, we have briefly introduced the proposed method which is based on copula function

### Modeling differential coexpression using Copula

Differential coexpression is simply defined as the change of coexpression patterns of a gene pair across two conditions. A straightforward method to measure this is to take the absolute difference of correlations between two gene expression profiles in two conditions. For a gene pair *gene*_*i*_ and *gene*_*j*_, this can be formally stated as: 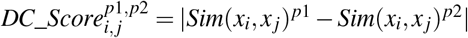, where *p*_*1*_, *p*_*2*_ are two different phenotype conditions, and *x*_*i*_, *x*_*j*_ represent expression profile of *gene*_*i*_ and *gene*_*j*_ respectively. Here *Sim*(*x*_*i*_, *x*_*j*_)^*p*^ signifies Pearson correlation between *x*_*i*_ and *x*_*j*_ for phenotype *p*.

In the statistical analysis, the simple way to measure the dependence between the correlated random variable is to use copulas^20^. Copula is extensively used in high dimensional data applications to obtain joint distributions from a random vector, easily by estimating their marginal functions.

Copulas can be described as a multivariate probability distribution function for which the marginal distribution of each variable is uniform. For a bivariate case copula is a function: *C*: [0, 1]^2^ → [0, 1], and can be defined as: *C*(*x*, *y*) = *P* (*X* ≤ *x*,*Y* ≤*y*), for 0 ≤ *x*,*y* ≤ 1, where *X* and *Y* are uniform random variable. Let, *Y*1 and *Y*2 be the random vectors whose marginals are uniformly distributed in [0, 1] and having marginal distribution *F*_*Y*1_ and *F*_*Y*2_ respectively. By Sklar’s Theorem^21^ we have the following: there exists a copula *C* such that *F*(*y*1, *y*2) = *C*(*F*_*Y1*_(*y*1), *F*_*Y2*_(*y*2)), for all *y*1 and *y*2 in the domain of *F_Y_* 1 and *F_Y_* 2. In other words, there exists a bivariate copula which represents the joint distribution as a function of its marginals. For the multivariate case the copula (C) function can be represented as:

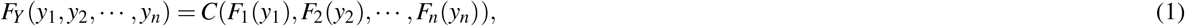

where (*Y*_1_, *Y*_2_,…,*Y*_*n*_) be the random vectors whose marginals are *F*_1_(*y*_1_), *F*_2_(*y*_2_),…, *F*_*n*_(*y*_*n*_). The converse of the theorem is also true. Any copula function with individual marginals *F_i_*(*y_i_*) as the arguments, represents valid joint distribution function. Assuming *F*(*Y*_1_, *Y*_2_,…,*Y*_*n*_) has *n*^*th*^ order partial derivatives, relation between the joint probability density function and the copula density function, say *c*, can be obtained as,

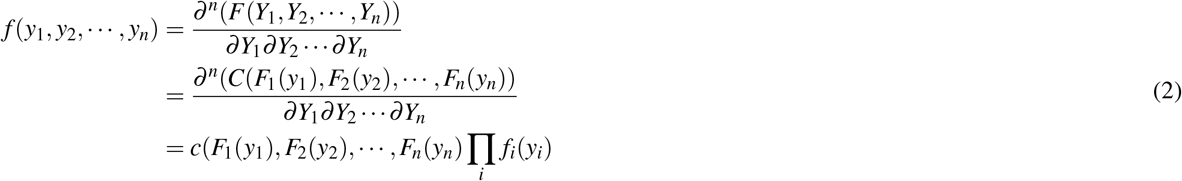

where, we define,

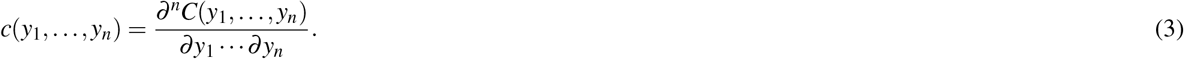

So, Copula is also known as joint distribution generating function with a separate choice of marginals. Hence, different families (parametric and non-parametric) of copulas exist which model different types of dependence structure. The example includes Farlie-Gumbel-Morgenstern family (parametric), Archimedean Copula (parametric), Empirical Copula (non-parametric), Gaussian (parametric), t (parametric) etc. Empirical copulas are governed by the empirical distribution functions, which tries to estimate the underlying probability distribution from given observations.

*Empirical copula* is defined as follows. Let *Y*_1_,*Y*_2_,…, *Y*_*n*_ be the random variables with marginals cumulative distribution function *F*_1_(*y*_1_), *F*_2_(*y*_2_),…, *F*_*n*_(*y*_*n*_), respectively.

The empirical estimate of (*F*_*i*_, *i* = 1,…, *n*), based on a sample, {*y*_*i*1_, *y*_*i*2_,…, *y*_*im*_} of size *m* is given by

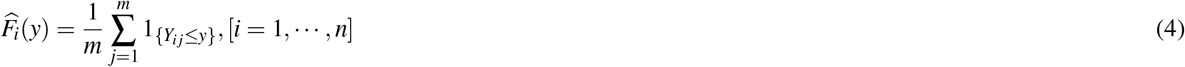

The *Empirical Copula* of *Y*_1_, *Y*_2_,…, *Y*_*n*_ is then defined as

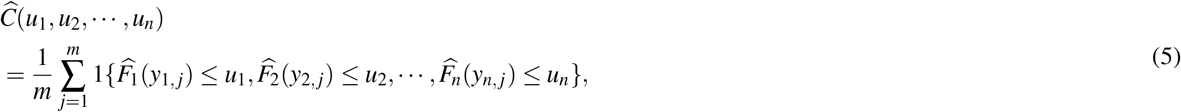

for *u*_*i*_ ∈ [0, 1], [*i* = 1,…, *n*].

Here, we model the dependence between each pair of gene expression profile using empirical copulas. As we were unaware of the distributions of expression profiles, so empirical copulas are the only choice here. Notably, we have estimated joint empirical copula density from the marginals of each gene expression profile. We have used beta kernel estimation to determine the copula density directly from the given data. The smoothing parameters are selected by minimizing the Asymptotic Mean integrated squared error (AMISE) using the Frank copula as the reference copula. The input to the copula density estimator is of size *n* × 2, where *n* is the number of samples in different datasets. For each pair of samples, we estimate the empirical copula density using beta kernel estimator. We have shown some marginal normal contour plots of copula density during the estimation process of BRCA dataset in the supplementary figure-1. To model the differential coexpression of a gene pair, we have measured a statistical distance between two joint distribution provided by the copulas. We have utilized the Kolmogorov-Smirnov (K-S) test to quantify the distance between two empirical distributions. Value of d-statistic represents the distance here. Thus, the distance obtained for a gene pair is treated as a differential coexpression score.

To check whether the distance between the joint distribution perfectly models the differential coexpression, we have performed an analysis. To show the concordance between the *DC*_*Score* with the proposed distance we have performed the following analysis. We create a 20 × 20 matrix *M*, whose rows (i) and columns (j) annotated with correlation values from −1 to +1 with 0.1 spacing. We create two pairs of marginals (*F*_*x*1_, *F*_*x*2_) and (*F*_*y*1_, *F*_*y*2_) having correlations *i* and *j* respectively. For the generation of marginals, we have used mvnorm function of MASS R-package and set the ‘empirical’ parameter of mvnorm as ‘TRUE’. This generates (*F*_*x*1_, *F*_*x*2_) or (*F*_*y*1_, *F*_*y*2_) with an exact correlation of *i* or *j*. Next, we compute joint distributions using copula function *F*_*X*1*X*2_ = *C*(*F*_*x*1_, *F*_*x*2_), *F*_*Y*1*Y*2_ = *C*(*F*_*y*1_, *F*_*y*2_) and finally compute KS distance between *F*_*X*1*X*2_ and *F*_*Y*1*Y*2_. Each entry of (i,j) in *M* is filled with this distance value. We generate *M* 100 times following the same method. Now to visualize the matrices each row is represented as a series of boxplot in the figure 1. For a fixed row, the *DC*_*Score* will increase from left to right along the column as it ranges from correlation value −1 to +1. Each facet in the figure corresponds a row/column in the matrix, which represents 20 sets of 100 distances corresponding to the correlations ranging from −1 to +1 with a spacing of 0.1. Considering each facet of the plot, it can be noticed that distances are gradually increasing with the increase in the *DC*_*Score*. For example, considering the second facet (corr value=-0.9), the distances increased from left to right gradually. So, it is evident from the figure that there exists a strong correlation between the distance and *DC*_*Score*, which signifies the proposed method can able to model the difference in coexpression patterns.

**Figure 1.**
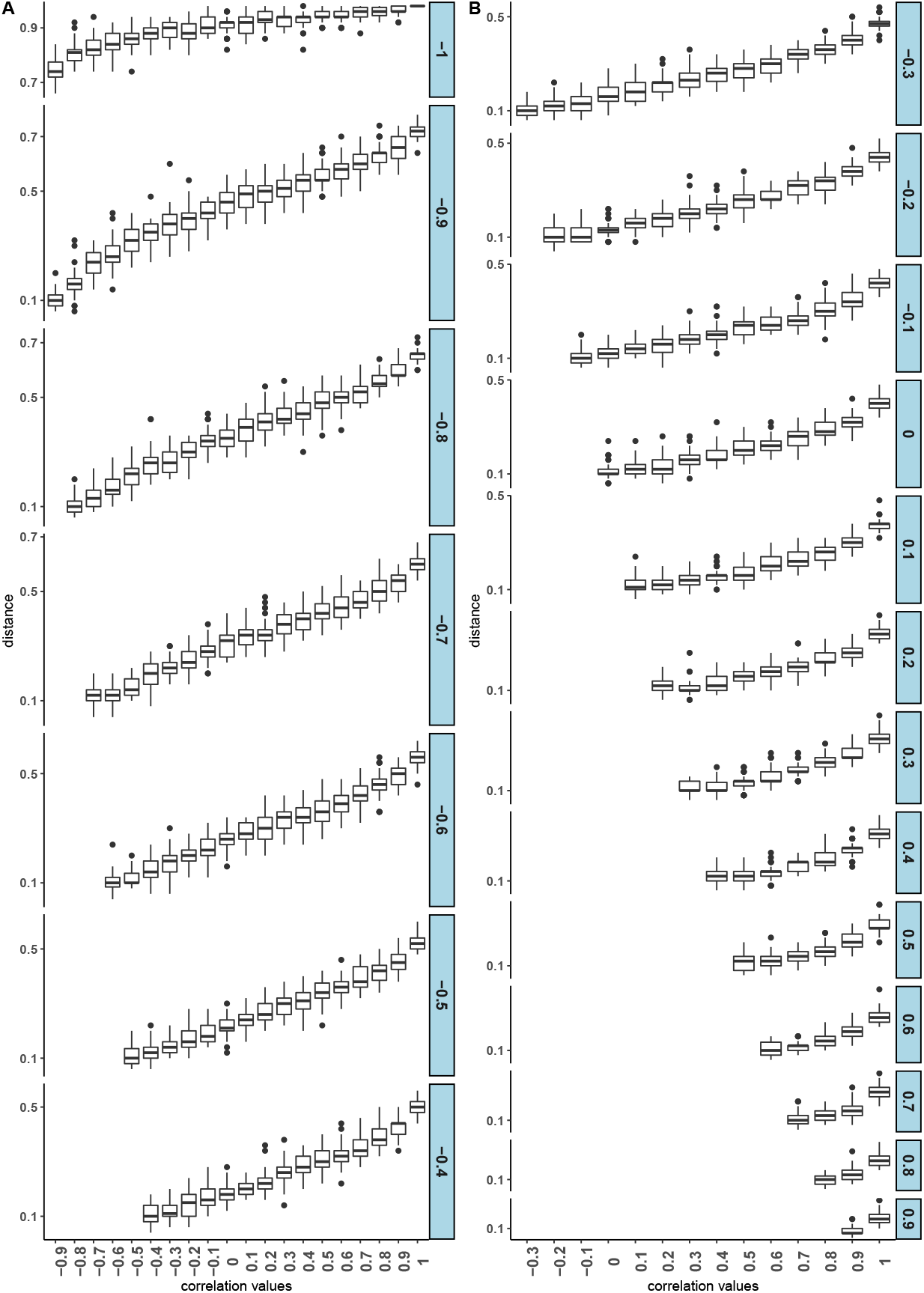
Boxplot showing the dependency between *DC*_*Score* and K-S distance, between two joint distributions. Panel-A shows the distances for the facets from correlation −1 to −0.4 and Panel-B shows the same for correlation −0.3 to +1.

### Stability of CODC

CODC is stable under noisy expression data. This is because of the popular “nonparametric”, “distribution-free” or “scale-invariant” nature of the copula^22^. The properties can be written as follows: Let, *C*_*XY*_ be a copula function of two random variables *X* and *Y*. Now, suppose *α* and *β* are two functions of *X* and *Y* respectively. The relation of *C*_(*α*(*X*),*β*(*Y*))_ and *C*_*XY*_ can be written as follows.

- **Property 1:** If *α* and *β* are strictly increasing functions, then the following is true:

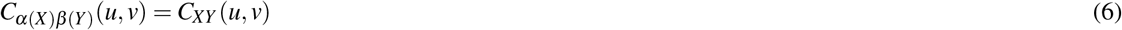
- **Property 2:** If *α* is strictly increasing and *β* is strictly decreasing, then the following holds:

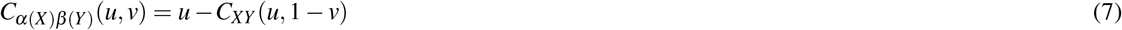
- **Property 3:** If *α* is strictly decreasing and *β* is strictly increasing function, then we have

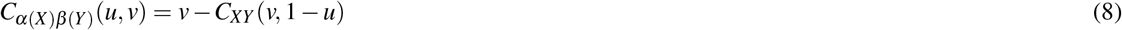
- **Property 4:** If *α* and *β* both are strictly decreasing function then the following holds:

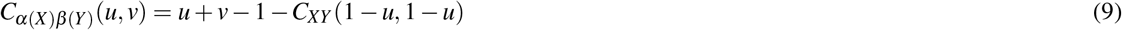

These properties of copula are used to prove that the distance measure used in CODC is approximately scaled invariant. Theoretical proves are described below and simulation result is given later in section **??**. The proof is as follows: we know the Kolmogorov-Smirnov statistic for a cumulative distribution function F(x) can be expressed as:

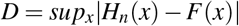

where *H*_*n*_ is an empirical distribution function for n i.i.d observation *X*_*i*_ ≤ *x*, and *sup* corresponds to supremum function. The two sample K-S test is used in CODC can be described similarly:

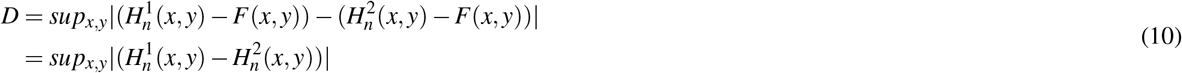

where 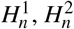 are denoted as the joint empirical distribution for two samples taken from normal and cancer respectively. Now the D-statistic can be written as:

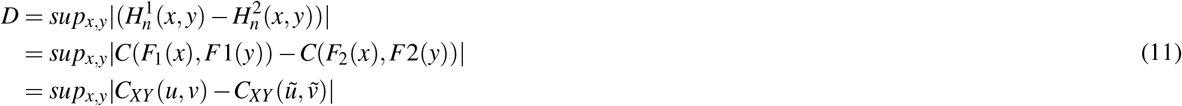

where C(.) is copula function and 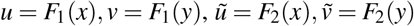 are uniform marginals of joint distribution 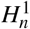 and 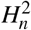

Let us assume that both *α* and *β* functions are strictly increasing. Then from equations 6 and 11 the distance *D* between 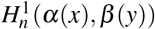 and 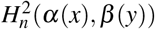 have the form

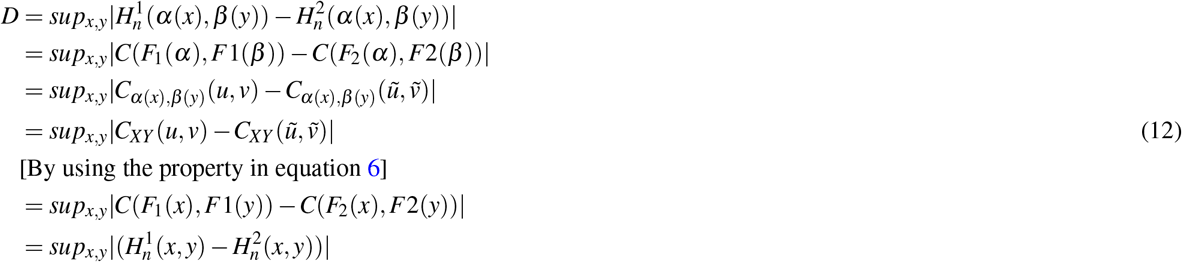

Now if *α* is strictly increasing and *β* is strictly decreasing then *D* can be written as:

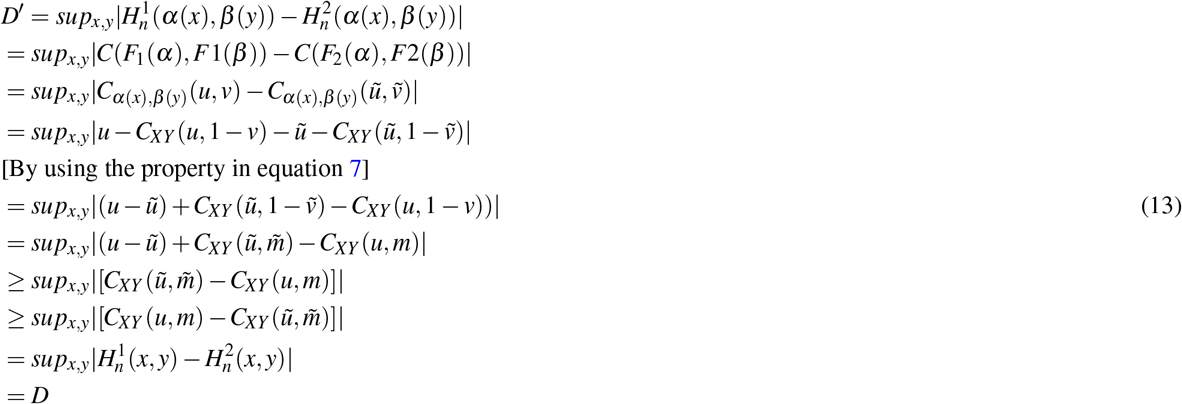

similarly, for strictly increasing *β* and strictly decreasing *α* the distance *D′* between 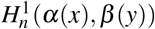 and 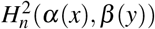 can be shown to satisfy the relation:

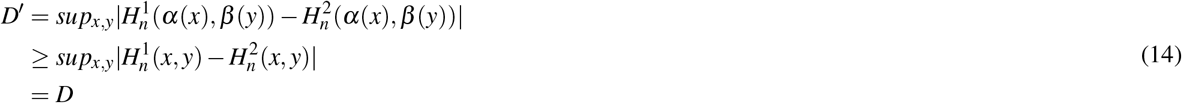

Finally let us consider *α* and *β* both are strictly decreasing function. The distance *D′* can be described as:

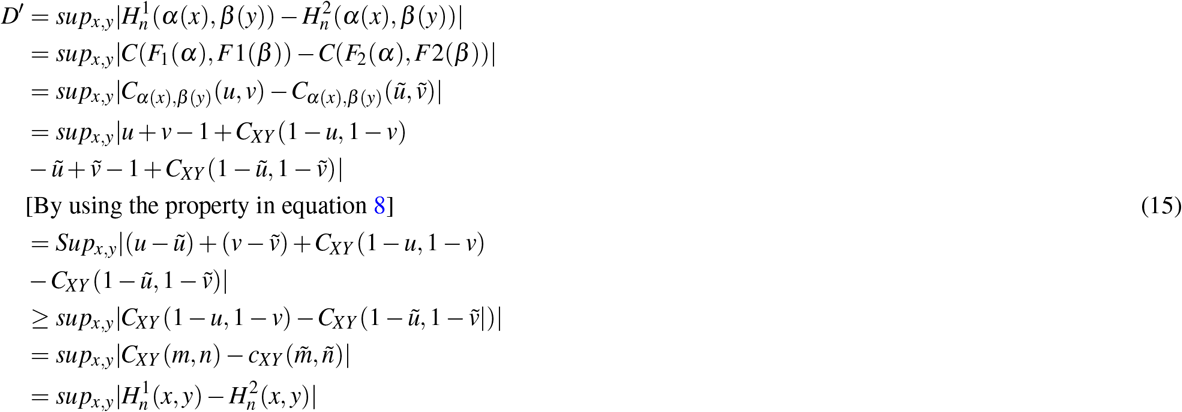

Thus the value of *D′* between two joint distribution 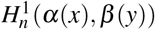 and 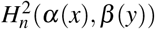 is the same as that of *D* which represents the distance 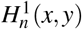 and 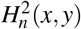 when both *α* and *β* are increasing function. For other cases of *α* and *β*, *D′* attains at least the value of *D*. So, the distance for two random variable *α*(*X*) and *β*(*Y*) is equal or at least that of the random variables *X* and *Y.* CODC treats the distance *D* as differential coexpression score, thus it remains the same (or at least equal) under any transformation of *X* and *Y*.

## Results

### Dataset preparation

We have evaluated the performance of the proposed method in five RNAseq expression data downloaded from TCGA data portal (https://www.cancer.gov/about-nci/organization/ccg/research/structural-genomics/tcga). We have downloaded matched pair of tumor and normal samples from five pan cancer data sets: Breast invasive carcinoma (BRCA, #samples=112), head and neck squamous cell carcinoma (HNSC, #samples = 41), liver hepatocellular carcinoma (LIHC, #samples = 50),thyroid carcinoma (THCA, #samples = 59) and Lung Adenocarcinoma (LUAD, #samples = 58). For preprocessing the dataset we first take those genes that have raw read count greater than two in at least four cells. The filtered data matrix is then normalized by dividing each UMI (Unique Molecular Identifiers) counts by the total UMI counts in each cell and subsequently, these scaled counts are multiplied by the median of the total UMI counts across cells^23^. Top 2000 most variable genes were selected based on their relative dispersion (variance/mean) with respect to the expected dispersion across genes with similar average expression. Transcriptional responses of the resulting genes were represented by the log2(fold-change) of gene expression levels from paired tumor and normal samples. A brief description of the datasets used in this article is summarized in table 1. Figure 2-panel(A) and Panel-B represent box and violin plot of average expression value of samples for each dataset.

**Table 1.**
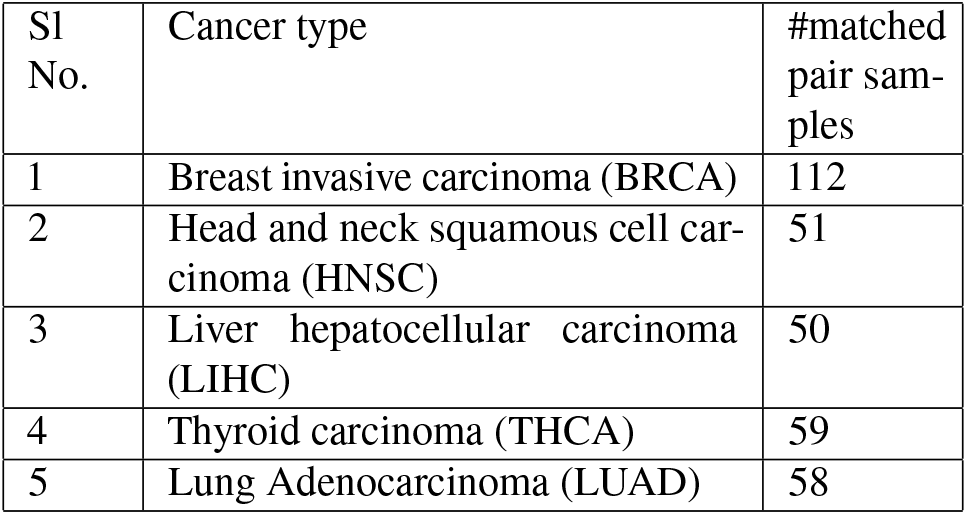
Tumor types and number of TCGA RNA-seq samples used in the analysis

| Sl No. | Cancer type | #matched pair samples |
| --- | --- | --- |
| 1 | Breast invasive carcinoma (BRCA) | 112 |
| 2 | Head and neck squamous cell carcinoma (HNSC) | 51 |
| 3 | Liver hepatocellular carcinoma (LIHC) | 50 |
| 4 | Thyroid carcinoma (THCA) | 59 |
| 5 | Lung Adenocarcinoma (LUAD) | 58 |

**Figure 2.**
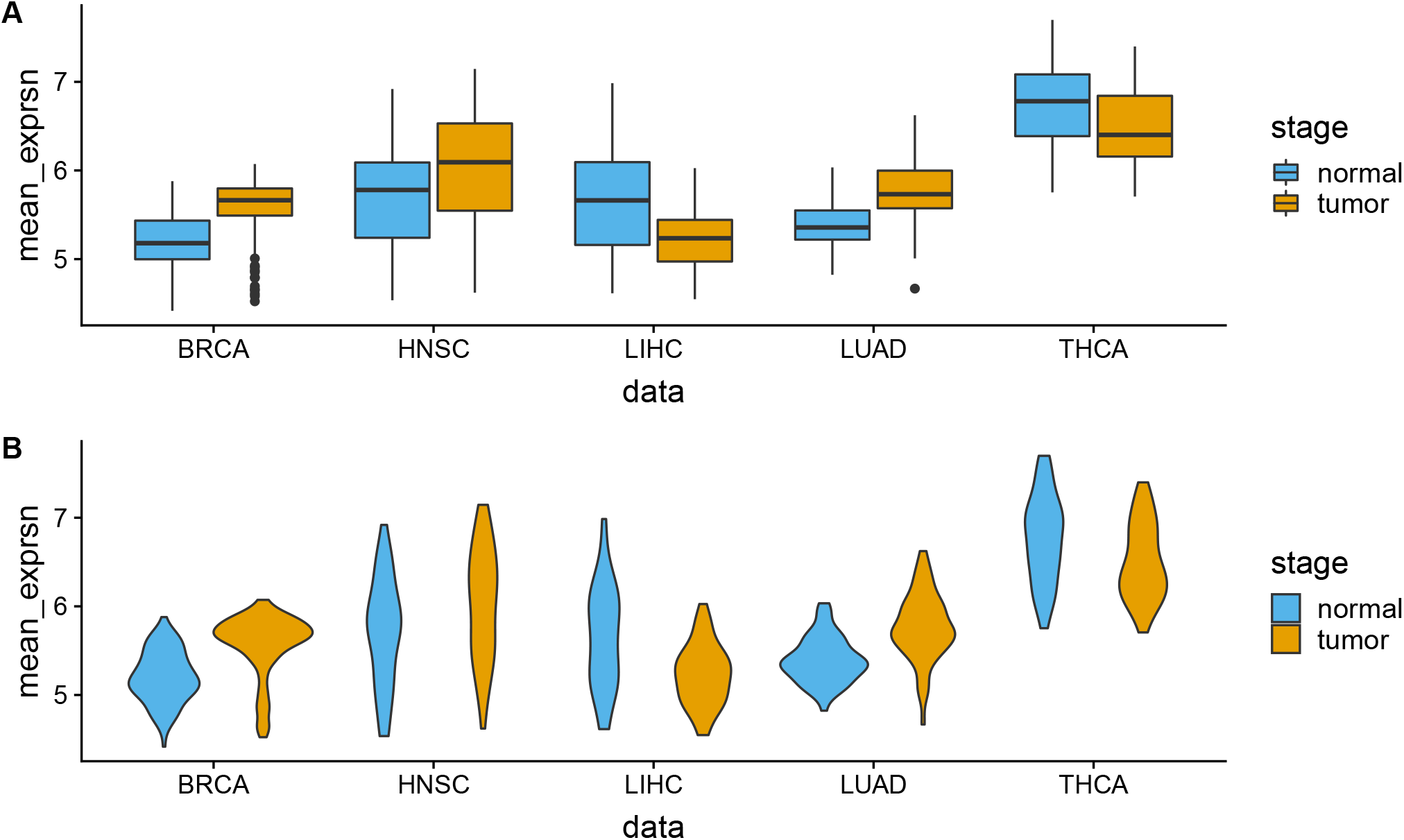
The figure describes box (panel-A) and violin plots (panel-B) of mean expression values of the used datasets.

### Detection of DC gene pair

Differential coexpression between a gene pair is modeled as a statistical distance between the joint distributions of their expression profiles in a paired sample. Joint distribution is computed by using empirical copula which takes expression profile of a gene as marginals in normal and tumor sample. The K-S distance, computed between the joint distribution is served as differential coexpression score between a gene pair. The score for a gene pair (*g_i_, *g*_*j*_*) can be formulated as: 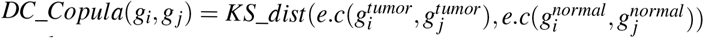, where KS-dist represents Kolmogorov-Smirnov distance between two joint probability distribution, e.c represents empirical copula, 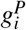 represents the expression profile of gene *g*_*i*_ at phenotype P. For each RNA-seq data, we have computed the *DC*_*Copula* matrix, from which we identify differentially coexpressed gene pairs.

To know how the magnitude of differential coexpression is changing with the score we plot the distribution of correlation values of gene pairs with their scores in Figure 3. The figure also shows the number of gene pairs having positive and negative correlations in each stage (normal/tumor). It can be noticed from the figure that high scores produce differentially coexpressed gene pairs having a higher positive and negative correlation. We collected the gene pairs having the score greater than 0.56 and plot the correlations values in Figure 4. This figure shows plots of all gene pairs having a positive correlation in normal and the negative correlation in tumor (shown in the panel-A) and vice-versa (shown in the panel-(B)). The density of the correlation values is shown in panel-C and panel-D for each case. In Figure 5 we create a visualization of top differentially coexpressed gene pairs in BRCA data which shows a strong positive correlation in tumor stage and negative correlation in normal stage. The Figure shows a heatmap of binary matrix constructed from the expression data of those gene pairs in tumor and normal stages. When the expression values showing the same pattern for a gene pair it is assumed 1, while 0 representing a non-matching pattern. From the Figure, it is quite understandable that most of the entry in the normal stage is 0 (non-match) while in tumor stage is 1 (match). For other datasets, the plots are shown in supplementary Figure-2.

**Figure 3.**
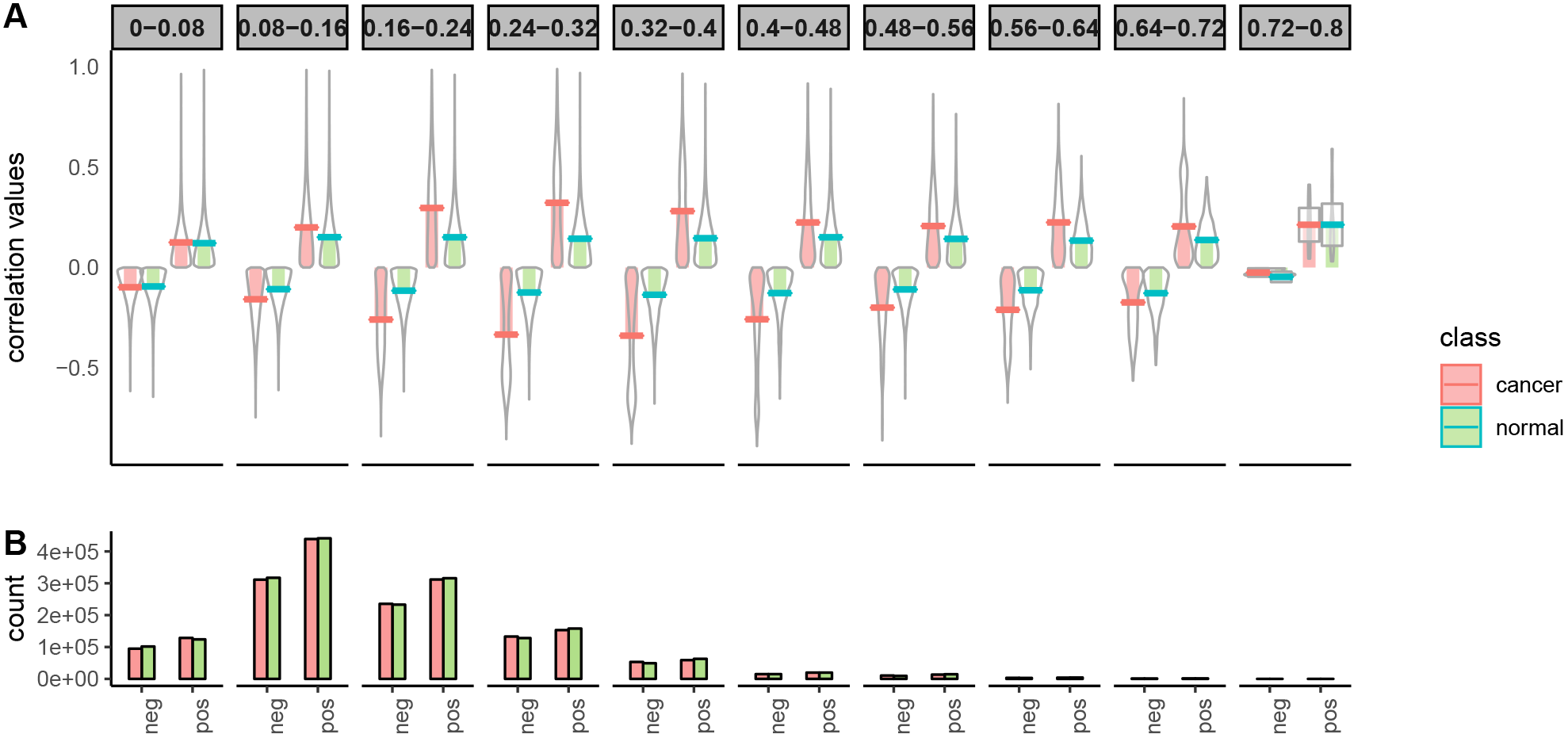
The Figure shows the distribution of correlation values in normal and cancer samples of BRCA data with the *DC*_*Copula* score. Panel-A shows the distribution for different *DC*_*Copula* scores. Here, four pirate plots are shown in each facet, two for positive and two for negative correlations. The violins in each facet represent the distribution of positive and negative correlations of gene pair in normal and cancer samples. Panel-B shows a bar plot representing the number of positive and negatively correlated gene pairs in normal and cancer samples in each facet.

**Figure 4.**
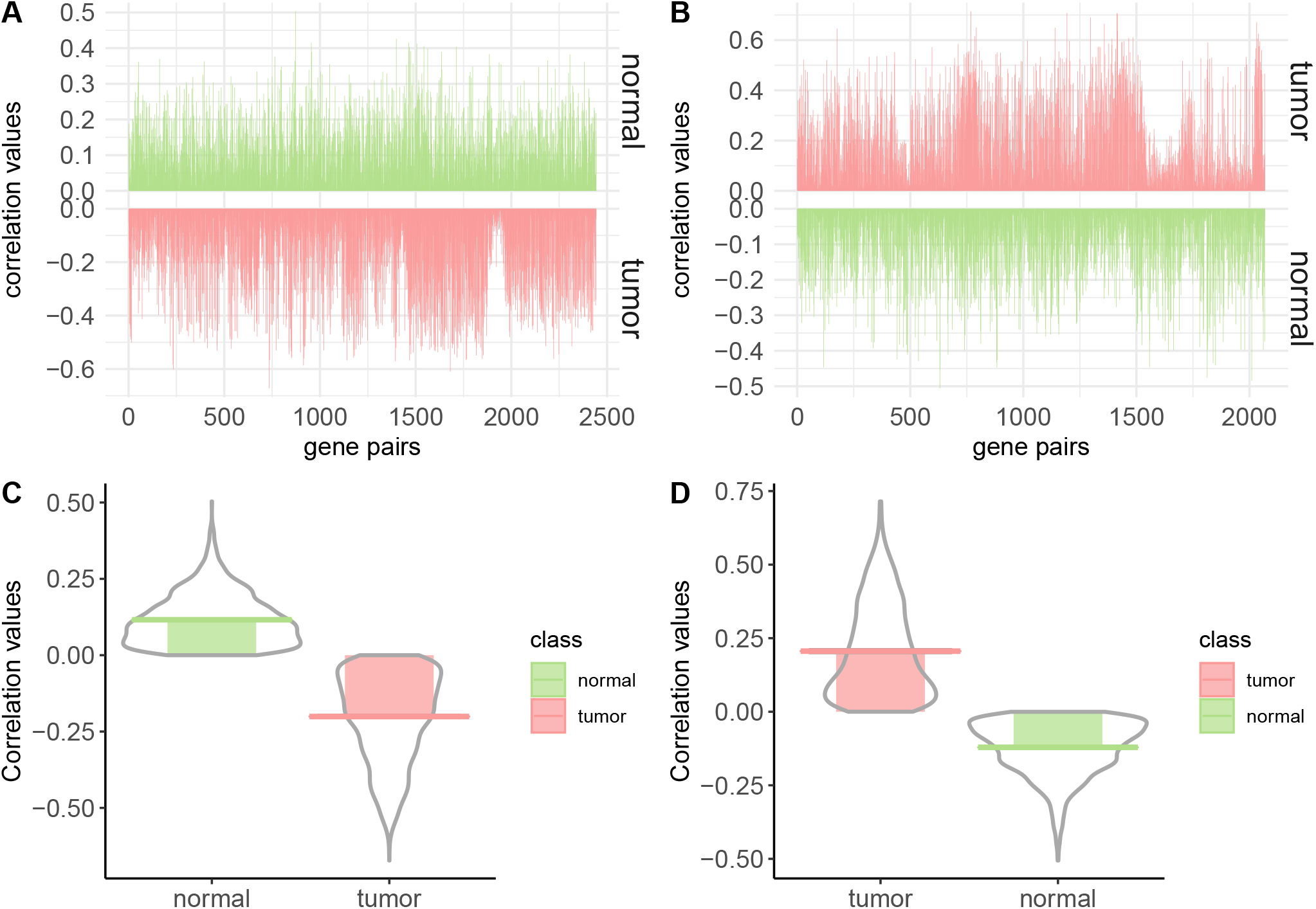
The Figure shows visualizations of gene pairs having *DC*_*copula* score greater than 0.56. Panel-A and Panel-B show the visualization of correlation values of gene pair having a positive correlation in normal and negative correlation in tumour and vice-versa, respectively. Panel-C and Panel-D represent the distribution of correlation values according to panel-A and panel-B, respectively.

**Figure 5.**
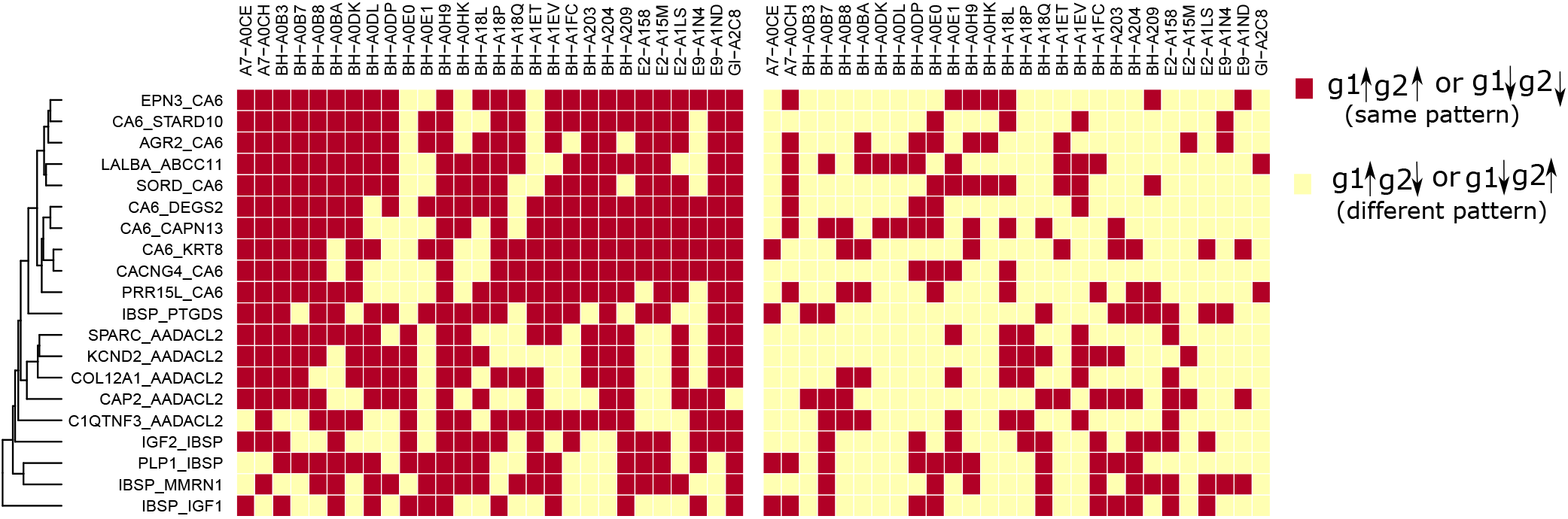
The Figure shows a heatmap representation of binary matrix constructed from the expression matrix of top differentially co-expressed gene pairs in normal and tumor stages. Expression values of a gene pair showing the same pattern are indicated as 1 and showing a different pattern is indicated as 0 in the matrix. The columns representing differentially co-expressed gene pairs while rows are the samples of BRCA data.

### Stability performance of CODC

To prove the stability of CODC we have performed the following analysis:

First, we add Gaussian noise to the original expression data of normal and cancer sample to transform these into noisy datasets. We use the rnorm function of R to create normally distributed noise with mean 0 and standard deviation 1 and we add this into the input data. We have utilized BRCA data for this analysis.

First, we compute the K-S distance and then obtain *DC*_*copula* matrix for both original and noisy datasets. Let us denote these two matrices as *D* and *D′*.

The usual way is to Pick a threshold *t* for *D* (or *D′*) and extract the gene pair (i,j) for which *D*(*i, j*)(or*D′*(*i, j*)) ≥ *t*. First, we set *t* as the maximum of *D* and *D′*, and then decreases it continuously to extract the gene pairs. For each *t*, we observe the number of common gene pairs obtained from *D* and *D′*. Figure 6 shows the proportion of common genes selected from *D* and *D′* for different threshold selection and different level of noise. Theoretically, CODC produces *D* with scores no more than *D′* (from sec). So, it is quite obvious that the number of common genes increased with a lower threshold value. From the property of section it can be noticed that the scores in *D* get preserved in *D′*. So, it is expected that obtained gene pairs from original data are also preserved in noisy data. Figure 6 shows the evidence for this case. As can be seen from the figure that even the noise label is 80%, for threshold value above 0.25 more than 55% of the gene-pairs are common between noisy and original datasets.

**Figure 6.**
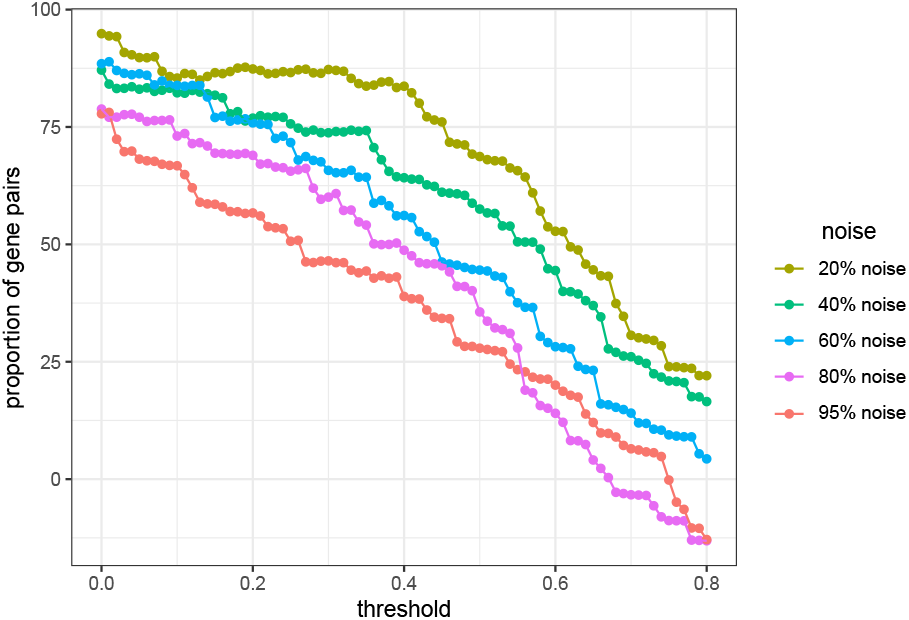
The proportion of common gene pairs obtained from noisy and original dataset with different threshold values and different noise level.

### Detection of differentially coexpressed modules

Detection of DC gene modules is performed by using hierarchical clustering on the DC matrix. Here, the differential coexpression score obtained from each gene pair is treated as the similarity measure between genes. The distance between a gene pair is formulated as: *dist*_*copula*(*g*_*i*_, *g*_*j*_) = 1 − *DC*_*copula*(*g*_*i*_, *g*_*j*_). For each dataset, modules are extracted using average linkage hierarchical clustering by using the *dist*_*copula* as a dissimilarity measure between a pair of a gene. For BRCA and HNSC data we have identified 15 modules, for LIHC data 14 modules, for LUAD 21 modules, and for THCA 22 modules are identified. For studying the relationship among the modules we have identified module eigengene networks for each dataset. According to^15^ module, eigengene represents a summary of the module expression profiles. Here, module eigengene network signifies coexpression relationship among the identified modules in two stages. We create visualizations of the module eigengene network for normal and tumor stages in figure 7. The upper triangular portion of the correlation matrix represents the correlation between module eigengenes for normal samples whereas the lower triangular portion represents the same for tumor samples. This figure shows the heatmap for BRCA, HNSC, and LUAD dataset. It is clear from figure 7 that most of the modules show differential coexpression pattern in normal and tumour stage. For a differentially coexpressed module, it is expected that it shows an opposite correlation pattern in two different phenotype conditions. Here, the correlation pattern between two modules is represented as the correlation between the module eigengenes. In the heatmap of Figure 7, we can observe that in all three datasets the correlation pattern between most of the MEs in normal and tumour stages has the opposite direction. For example, from panel-A, it can be noticed that for BRCA data, the modules have a negative correlation in normal phase while showing a strong positive correlation in tumour phase. For HNSC dataset, the opposite case is observed. Modules have a strong positive correlation in normal phases while having a negative correlation in tumor phase. In supplementary figure-2, the visualization of all datasets is given.

**Figure 7.**
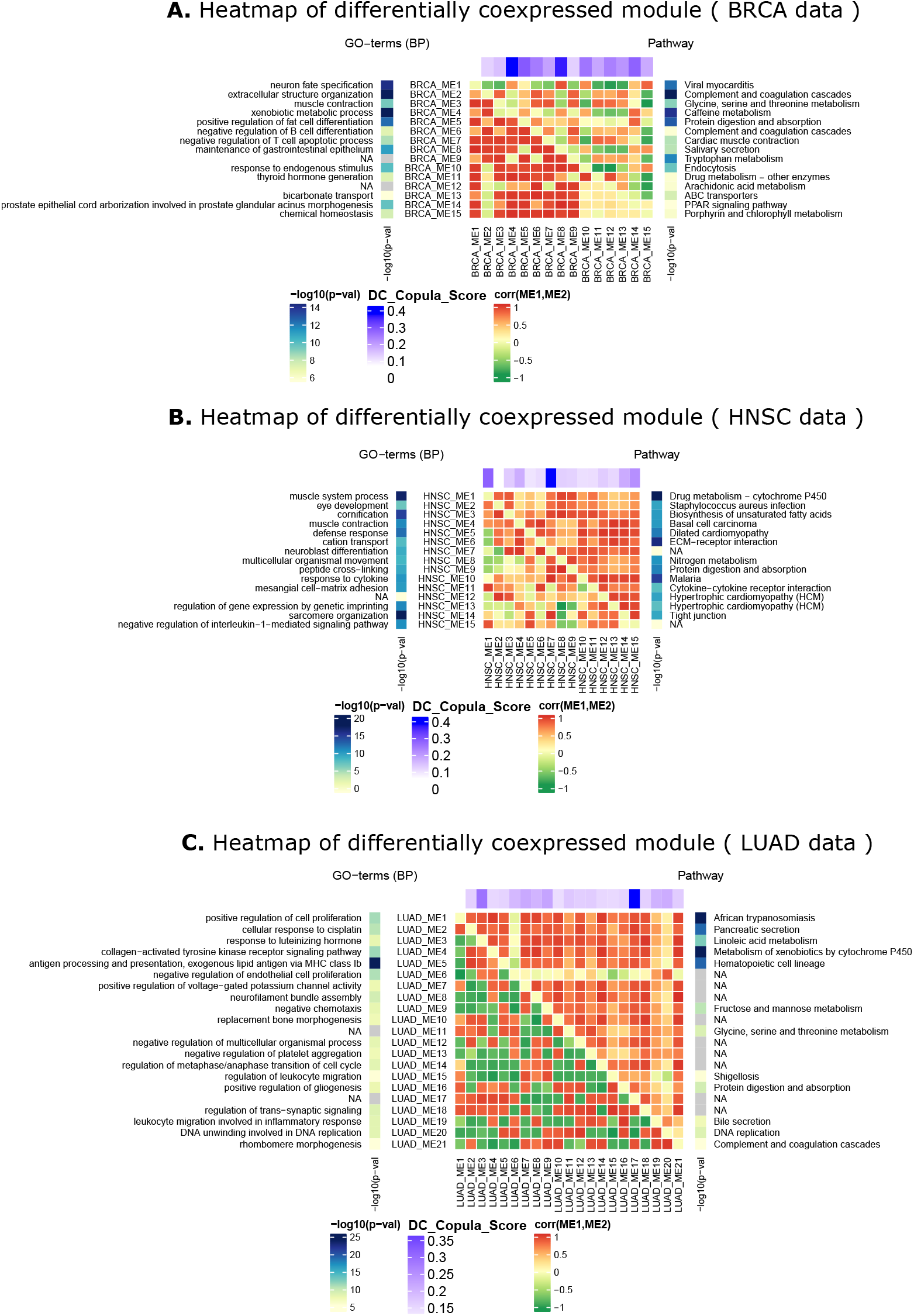
Heatmap of differentially coexpressed modules. Here the heatmap is shown for module eigengenes.The upper triangular portion of the matrix represents correlations of module eigengenes in normal samples whereas lower triangular portion signifies the same for tumor samples. Left and right sidebar of the heatmap represents −log(p-value) of significantly enriched GO-terms and pathway, respectively. ‘NA’ stands for unavailability of significant pathway or GO terms. Upper annotation bar of the heatmap shows the *DC*_*copula* score of the module. Panel-A shows the heatmap for BRCA data, whereas panel-B and Panel-C show heatmap HNSC and LUAD data.

### Comparisons with competing methods

For comparison purpose, we have taken three competing techniques such as Diffcoex, coXpress, and DiffCoMO and compared them with our proposed method. All these methods are extant DC based, which look for gene modules with altered coexpression between two classes. DiffCoEx performed hierarchical clustering on the distance matrix complied from correlation matrices of two phenotype stages. CoXpress detect correlation module in one stage and find the alternation of the correlation pattern within the module in other class. DiffCoMO uses the multiobjective technique to detect differential coexpression modules between two phenotype stages. We have made two approaches for comparing our proposed method with state-of-the art. We first compare the efficacy of these methods for detecting differential coexpressed gene pairs and next compare the modules identified in each case. For the first case, we take top 1000 gene pairs having high *DC*_*copula* scores from the DC matrix, and perform classification using normal and tumor samples. Expression ratio of each DC gene pairs from the expression matrix was taken and compiled a *n* × 1000, where *n* represents the number of samples in each data. For the other three methods, we have also selected the same number of differentially coexpressed gene pairs for the classification task. Table 2 shows some parameters we have used for the selection of the gene pairs. For CoXpress, first, we have used ‘cluster.gene’ and ‘cuttree’ function with default parameters provided in the R package of *CoXpress* to get the gene clusters according to the similarity of their expression profiles. These groups are then examined by the *coXpress* R function to identify the differentially coexpressed modules by comparing with the *t*-statistics generated by randomly resampling the dataset 10,000 times for each group. We have taken top 10 modules based on the robustness parameter, which tells the number of times that the group was differentially co-expressed in 1000 randomly resampled data. Now we have selected 1000 gene pairs randomly from those modules. For DiffCoEx method, we collected the DC gene pairs before partitioning them in modules. We used the code available in the supplementary file of the original paper of DiffCoEx, to get the distance score matrix which is used in the hierarchical clustering for module detection. We sort the score of the distance matrix and pick top 1000 gene pairs based on the scores. For DiffCoMO we use the default parameters to cluster the network to obtained differentially coexpressed modules. As it utilized multiobjective method, so all the Pareto optimal solutions of the final generation is taken as selected modules. We then choose 1000 gene pairs randomly from the identified modules. Classification is performed by treating normal and tumor samples as class label. A toy example of the comparison is shown in figure 8. Please note that all these methods are meant for differentially coexpressed module detection. So, for comparison, we collected the DC gene pairs before partitioning them in modules. We train four classifiers Boosted GLM, Naive Bayes, Random Forest and SVM with the data and take the classification accuracy. The classification results are shown in the figure 9. It can be noticed from the figure that for most of the dataset proposed method achieved high accuracy compare to the other methods.

**Table 2.**
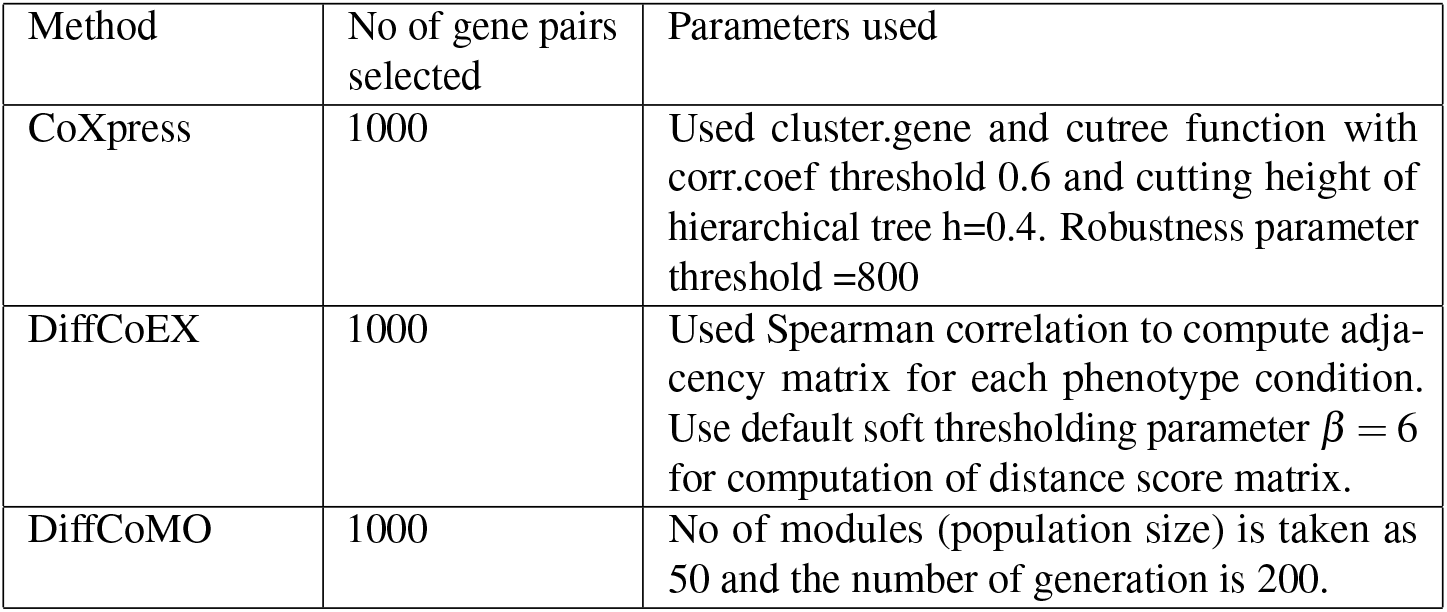
table shows the different parameters/threshold we have used for selecting differentially coexpressed gene pairs for other methods

**Figure 8.**
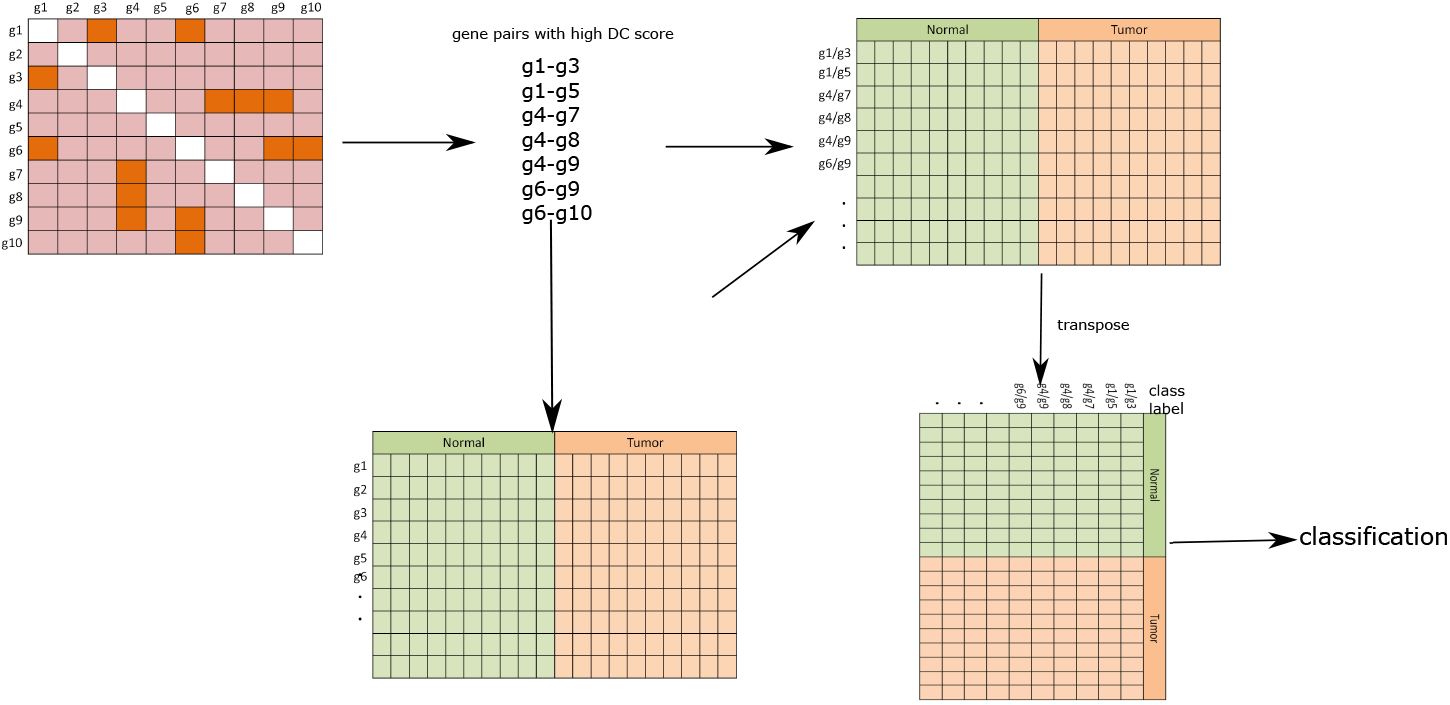
A toy example of performing classification on differentially coexpressed gene pairs. From the DC matrix top gene pairs are selected based on *DC*_*copula* score. Expression ratio is computed for each gene pairs for normal and tumor samples. The final matrix is then transposed and subsequently, classification is performed using normal and tumor sample as class label.

**Figure 9.**
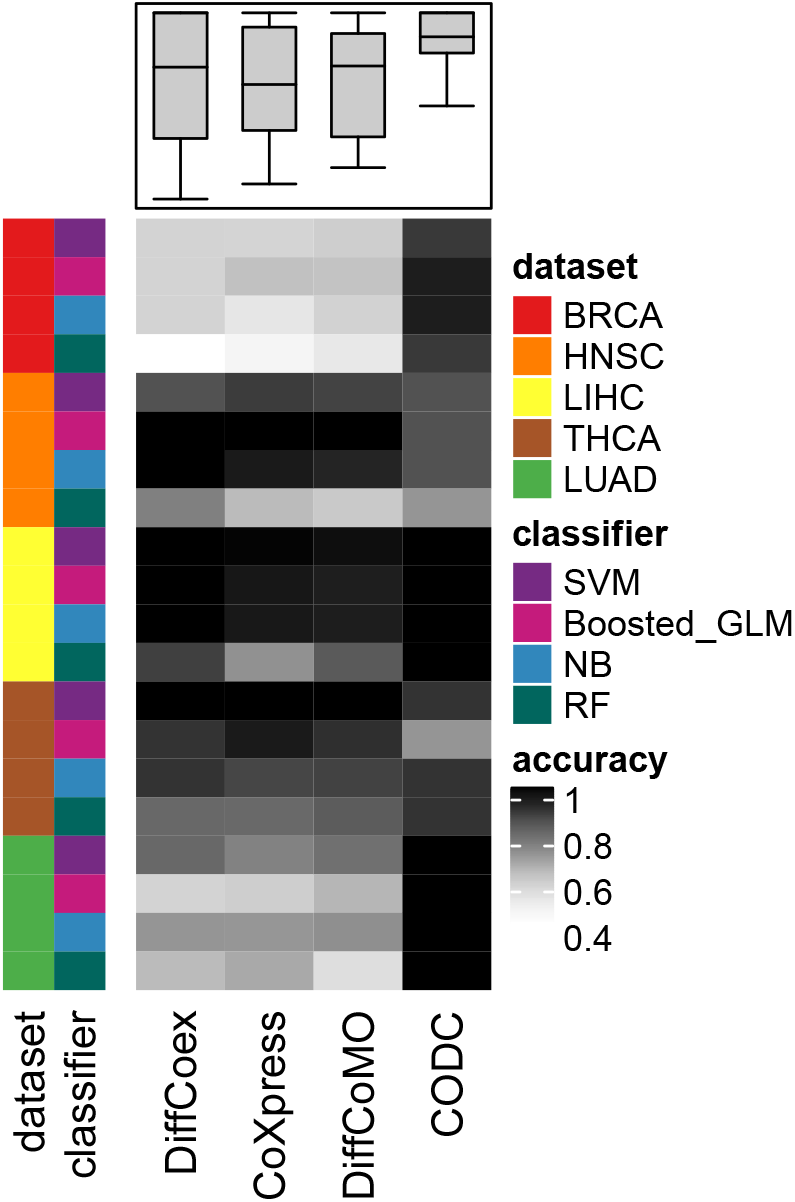
Comparison of classification accuracy for five datasets with four classifiers BGM, Naive-Bayes, Random Forest and SVM.

To assess the performance of all the methods for detecting differential coexpression modules, we check the distribution of correlation score of gene pairs within top modules in normal and tumor samples. Extant methods do a comparison by computing the absolute change in correlation value between a pair of a gene within a module. The problem for this type of comparison is that the score ignores a small change in differentially coexpression. It also fails to consider the gene pair having a low score but and correlation of opposite sign in two conditions. For example, it emphasized the gene pair with correlation value 0.2 in normal and 0.7 in the tumor (here the score is 0.5) rather than the gene pair whose correlation value is −0.2 in normal and 0.2 in the tumor (here the score is 0.4). So, for comparison, it is required to investigate the number of gene pairs having correlation values of an opposite sign over −1 to +1. So, for all identified modules we calculate the correlation score of each gene pairs in two different samples (normal and cancer) and plot frequency polygon in figure 10. To investigate whether the gene pairs within the modules show a good balance in positive and negative correlations, we have computed the correlation score for all the identified modules of DiffCoMO, DiffCoEx, and CoXpress. Figure 10 shows the comparisons of the correlation scores. It is noticed from the figure that gene pairs within the identified modules of the proposed method show good balance in positive and negative correlation values. DiffCoMO and DiffCoEX have also achieved the same, whereas most of the gene pairs within the coXpress modules shifted towards positive correlation in both tumor and normal samples. In figure 10-(b) we have also shown the boxplot of the correlation values obtained from different methods. As can be seen from the figure, the median line of correlation values for the proposed method is nearer to 0, which signifies good distribution of correlation scores in normal and tumor samples over −1 to +1. Thus the proposed method can able to detect differentially coexpressed gene pairs having correlation values well distributed between −1 to +1.

**Figure 10.**
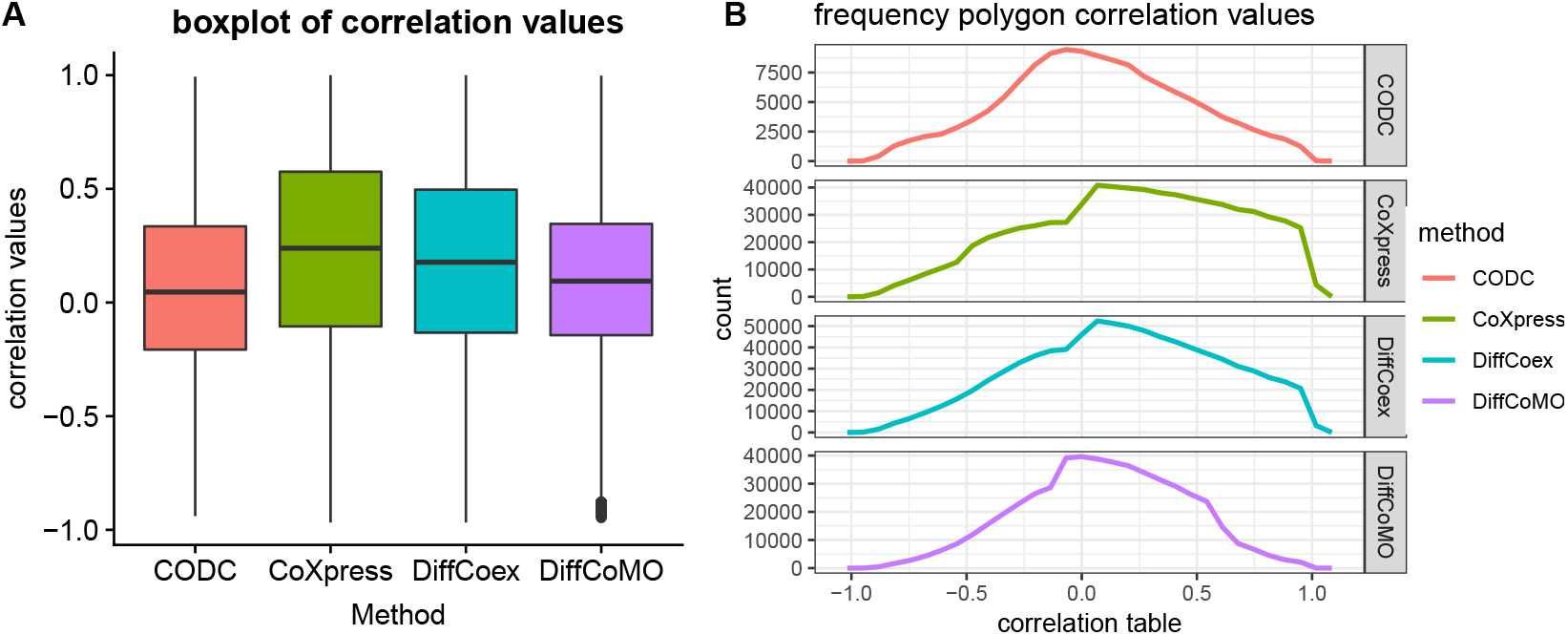
Distribution of correlation scores of the gene pairs in normal and tumor stage. Each facet shows the distribution for different methods.

To compare CODC with the other methods we tested its performance in a simulated dataset also. To create the simulated data we have used HNSC RNA-seq expression dataset. We create the simulated data as follows:

1. For each gene *g*_*i*_ in the normal sample, we simulated expression profile of sample *s*_*j*_ as 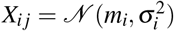 where *m*_*i*_ represents mean expression value of gene *g*_*i*_ across all 51 HNSC normal samples, and *σ*^2^ represents their variance.
2. Similarly we simulated expression profile of tumor sample 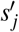 as 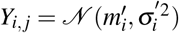, where 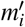 is mean of the expression value of gene 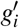 and 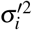 is the variance. Here, we assume that a gene pair is true differentially coexpressed if the following condition is hold: *DC*_*score*(*g*_*i*_, *g*_*j*_) > 0.5 and the correlation between *g*_*i*_ and *g*_*j*_ has opposite sign in tumor and normal stage.
3. We then add different levels of Gaussian noise to the *Y*_*ij*_ to create different noisy expression data 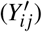 from the simulated tumor samples. We use rnorm function of R to produce normally distributed noise with mean 0 and standard deviation 1.

Now we compute differentially coexpressed gene pair between simulated normal (*X*_*ij*_) and noisy sample 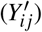 and compare them with the underlying true differentially coexpressed gene pairs. We compute the proportion of matched gene pairs and plot the results against all the different noise level in figure 12. We have done this analysis for all competing methods. To select the differentially coexpressed gene pairs from simulated normal and noisy simulated tumor sample we use *DC*_*Copula* threshold as 0.6. For other competing methods we use the threshold and other parameters same as previous analysis which are provided in table 2.

### Pathway analysis

To compare functional enrichment of identified modules we have utilized KEGG pathway enrichment analysis. We defined enrichment score of a pathway in a module as: 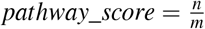, where n is the fraction of the pathway genes in the module and m is the fraction of the pathway genes in the dataset. We compare the pathway score for the modules identified for DiffCoEx, DiffCoMO, CoXpress and the proposed method.For comparison purpose we have utilized HNSC dataset. For identifying modules in other competing methods we have utilized the the parameters provided in table 2. For coXpress we take the modules with robustness parameter value greater than 760 which produces 19 clusters. For DiffcoEx we have used the default parameters for cutreeDynamic function (cutHeight=.996, minClusterSize = 20) which produces 42 clusters. Figure 11 shows the result. The Y-axis represents CCDF (complementary cumulative distribution function) which represents how often the number of modules is above a certain value of pathway score. It is clear from the figure that more modules for the proposed method achieved high pathway score compare to other competing methods. In figure 7 we have shown heatmaps of differentially coexpressed modules for BRCA, HNSC and for LUAD data. The heatmap also provides pathways and GO-terms significantly enriched with the modules. The p-value for KEGG pathway enrichment and GO enrichment is computed by using the hypergeometric test with 0.05 FDR corrections. We have utilized GOstats, kegg.db and GO.db R package for that. It can be seen from figure 7, panel-A that some pathways such as ‘Complement and coagulation cascades’,‘Proximal tubule bicarbonate reclamation’,‘Caffeine metabolism’, ‘Protein digestion and absorption’, ‘Tryptophan metabolism’ and ‘ABC transporters’, are strongly associated with the identified modules of BRCA. ‘Tryptophan metabolism’ have eminent evidence to linked with malignant progression in breast cancer^24^. In^25^ the association between ABC transporters with breast carcinoma has been established. From panel-B it can be seen that Drug metabolism-cytochrome P450^26^ ECM-receptor interaction^27^, ‘Nitrogen metabolism’, ‘Protein digestion and absorption’, are significantly associated with the modules of HNSC data. Some pathways such as ‘Drug metabolism-cytochrome P450’ and ‘ECM-receptor interaction’ have strong evidence associated with the Head and Neck Squamous Cell Carcinomas^26,27^. Similarly from panel-C it can notice that pathways such as ‘Metabolism of xenobiotics by cytochrome P450’, ‘Pancreatic secretion’, ‘Linoleic acid metabolism’ is significantly associated with modules of LUAD data. Among them there exist strong evidence for pathways: ‘Metabolism of xenobiotics by cytochrome P450’,^28^, ‘Pancreatic secretion’^29^, and ‘Linoleic acid metabolism’^30^ to be associated with lung carcinoma.

**Figure 11.**
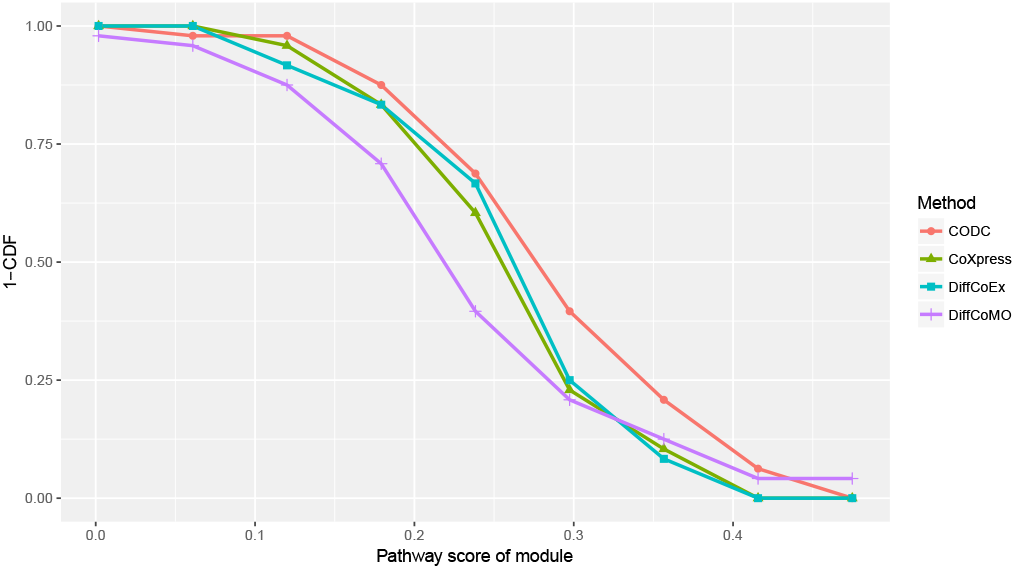
Distribution of pathway score for each of the comparing method. The figure shows the fraction of identified modules having a pathway score above a certain value.

**Figure 12.**
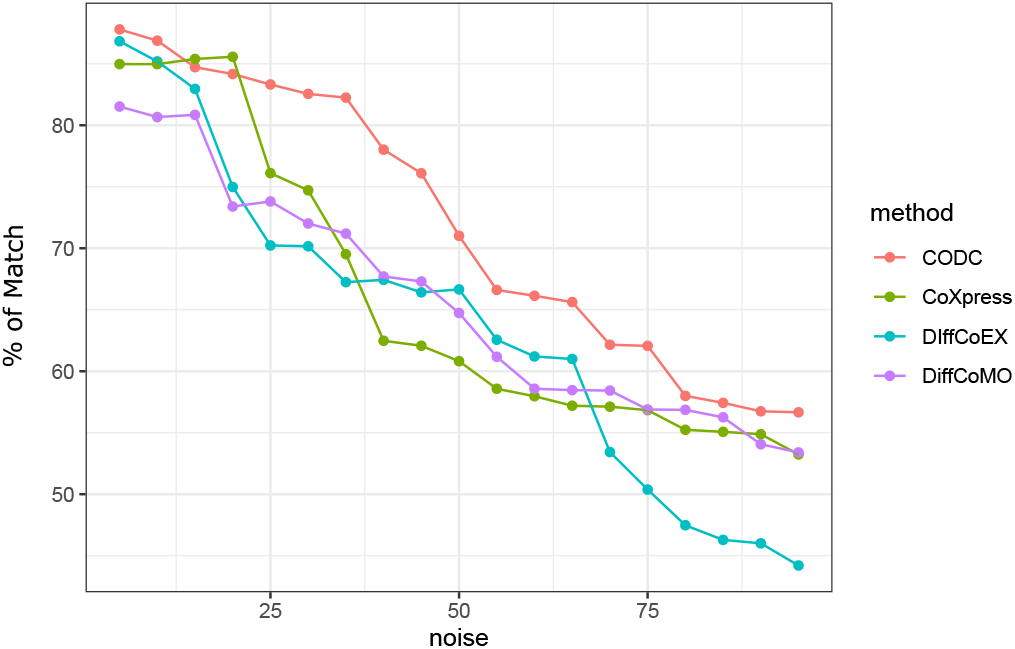
Figure shows percentage of matching between identified gene pairs with true differentially coexpressed gene pairs of simulated data. Results are shown for different noise level and for different method.

## Conclusion

In this article, we have proposed CODC, a copula based model to detect differential coexpression of genes in two different samples. CODC seeks to identify the dependency between expression patterns of a gene pair in two conditions separately. The Copula is used to model the dependency in the form of two joint distributions. Kolmogorov-Smirnov distance between two joint distributions is treated as differential coexpression score of a gene pair. We have compared CODC with three competing methods DiffCoex, CoXpress and DiffCoMO in five pan-cancer RNA-Seq data of TCGA. CODC’s ability for delineating minor change of coexpression in two different samples makes it unique and suitable for differential coexpression analysis. The scale-invariant property of copula inherited into CODC to make it robust against noisy expression data. It is advantageous for detecting the minor change in correlation across two different conditions which is the most desirable feature of any differential coexpression analysis.

Under the premise that the differential coexpressed genes are likely to be important bio-markers, we demonstrate that CODC identifies those which achieve better accuracy for classifying samples. Moreover, CODC goes a step further from the pairwise analysis of genes and seeks modules wherein differential coexpression are prevalent among each pair of genes. We have also analyzed the identified modules enriched with different biological pathways and highlighted some of these such as: ‘Complement and coagulation cascades’, ‘Tryptophan metabolism’, ‘Drug metabolism-cytochrome P450’, ‘ECM-receptor interaction’.

We have evaluated the efficacy of CODC on 5 different pan-cancer dataset to effectively extract differential coexpression gene pairs. Besides that, we have also compared different methods for detecting differentially coexpressed modules in those data. It is worth mentioning that CODC improves upon the competing methods. We have also proved that the scale-invariant property of copula makes CODC more robust for detecting differential coexpression in noisy data. The most important part of the DC analysis is to reveal changes in gene correlation that would not be detected by traditional DE analysis. As CODC uses copula for measuring gene-gene dependency and copula is a multivariate measure so it can be easily extendible to use in the measurement of dependence structure of multiple genes.

## Acknowledgment

We would like to acknowledge support from J.C. Bose Fellowship [SB/S1/JCB-033/2016 to S.B.] by the DST, Govt. of India; SyMeC Project grant [BT/Med-II/NIBMG/SyMeC/2014/Vol. II] given to the Indian Statistical Institute by the Department of Biotechnology (DBT), Govt. of India.

## Code Availability

https://github.com/Snehalikalall/CODC

## Data Availability

Datasets are available in TCGA data portal: (https://www.cancer.gov/about-nci/organization/ccg/research/structural-genomics/tcga)

## References

1. Ralston, A. & Shaw, K. Gene expression regulates cell differentiation. Nat. education 1, 127 (2008).

2. Yang, Y. et al. Gene co-expression network analysis reveals common system-level properties of prognostic genes across cancer types. Nat. communications 5, 3231 (2014).

3. Eisen, M. B., Spellman, P. T., Brown, P. O. & Botstein, D. Cluster analysis and display of genome-wide expression patterns. Proc. Natl. Acad. Sci. 95, 14863–14868 (1998).

4. Ideker, T. & Krogan, N. Differential network biology. Mol. Syst. Biol. 565 (2011).

5. Ray, S. & Bandyopadhyay, S. Discovering condition specific topological pattern changes in coexpression network: an application to hiv-1 progression. IEEE/ACM Transactions on Comput. Biol. Bioinforma. 11 (2015).

6. Cho, S., Kim, J. & Kim, J. Identifying set-wise differential co-expression in gene expression microarray data. BMC Bioinforma. 10 (2009).

7. Kostka, D. & Spang, R. Finding disease specific alterations in the co-expression of genes. Bioinformatics 20, i194–i199 (2004).

8. Lai, Y., Wu, B., Chen, L. & Zhao, H. A statistical method for identifying differential gene–gene co-expression patterns. Bioinformatics 20, 3146–3155 (2004).

9. Kostka, D. & R, R. S. Finding disease specific alterations in the co-expression of genes. Bioinformatics 20, i194–199. (2005).

10. Watson, M. Coxpress: differential co-expression in gene expression data. BMC Bioinforma. 7 (2006).

11. Tesson, B., Breitling, R. & Jansen, R. Diffcoex: a simple and sensitive method to find differentially coexpressed gene modules. BMC Bioinforma. 11 (2010).

12. Fang, G., Kuang, R., Pandey, G., Steinbach, M. & et. al, C. M. Subspace differential coexpression analysis: problem definition and a general approach. 145–156 (Pacific Symposium on Biocomputing, 2010).

13. Wu, G. & Stein, L. A network module-based method for identifying cancer prognostic signatures. Genome Biol. 13, DOI: 10.1186/gb-2012-13-12-r112 (2012).

14. Watson, M. Coxpress: differential co-expression in gene expression data. BMC Bioinforma. 7 (2006).

15. Langfelder, P. & Horvath, S. Wgcna: an r package for weighted correlation network analysis. BMC Bioinforma. 9 (2008).

16. Amar, D., Safer, H. & Shamir, R. Dissection of regulatory networks that are altered in disease via differential co-expression. Plos Comp Bio. 9, e1002955 (2013).

17. Ray, S. & Maulik, U. Identifying differentially coexpressed module during hiv disease progression: A multiobjective approach. Sci. reports 7, 86 (2017).

18. Nelsen, R. B. An introduction to copulas (Springer Science & Business Media, 2007).

19. Embrechts, P. Copulas: A personal view. J. Risk Insur. 76, 639–650 (2009).

20. Nelsen, R. B. Introduction. In An Introduction to Copulas, 1–4 (Springer, 1999).

21. Sklar, A. Random variables, joint distribution functions, and copulas. Kybernetika 9, 449–460 (1973).

22. Nelsen, R. B. Properties and applications of copulas: A brief survey. In Proceedings of the first brazilian conference on statistical modeling in insurance and finance, 10–28 (Citeseer, 2003).

23. Zheng, G. X. et al. Massively parallel digital transcriptional profiling of single cells. Nat. communications 8, 14049 (2017).

24. Juhász, C. et al. Tryptophan metabolism in breast cancers: molecular imaging and immunohistochemistry studies. Nucl. medicine biology 39, 926–932 (2012).

25. Hashimoto, K. et al. Activated pi3k/akt and mapk pathways are potential good prognostic markers in node-positive, triple-negative breast cancer. Annals oncology 25, 1973–1979 (2014).

26. Shatalova, E. G., Klein-Szanto, A. J., Devarajan, K., Cukierman, E. & Clapper, M. L. Estrogen and cytochrome p450 1b1 contribute to both early-and late-stage head and neck carcinogenesis. Cancer Prev. Res. 4, 107–115 (2011).

27. Kuang, J., Zhao, M., Li, H., Dang, W. & Li, W. Identification of potential therapeutic target genes and mechanisms in head and neck squamous cell carcinoma by bioinformatics analysis. Oncol. letters 11, 3009–3014 (2016).

28. Anttila, S., Raunio, H. & Hakkola, J. Cytochrome p450–mediated pulmonary metabolism of carcinogens: regulation and cross-talk in lung carcinogenesis. Am. journal respiratory cell molecular biology 44, 583–590 (2011).

29. Gonlugur, U., Mirici, A. & Karaayvaz, M. Pancreatic involvement in small cell lung cancer. Radiol. oncology 48, 11–19 (2014).

30. Barhoumi, R., Mouneimne, Y., Chapkin, R. S. & Burghardt, R. C. Effects of fatty acids on benzo [a] pyrene uptake and metabolism in human lung adenocarcinoma a549 cells. PloS one 9, e90908 (2014).

